# Genomic Analysis Reveals Genetic Diversity, Population Structure and Evolutionary Dynamics in Bluebunch wheatgrass (*Pseudoroegneria* species)

**DOI:** 10.1101/2025.03.03.641294

**Authors:** Yuanyuan Ji, Nadeem Khan, Raju Chaudhary, Sampath Perumal, Zhengping Wang, Pierre Hucl, Bill Biligetu, Andrew G. Sharpe, Lingling Jin

## Abstract

Bluebunch wheatgrass (BBWG: *Pseudoroegneria* species) is an outcrossing perennial grass of considerable ecological and agricultural importance due to its resilience in adverse environmental conditions. To gain a deeper understanding of the diversity, population structure, and genetic relationship within the *Pseudoroegneria* genus, we analyzed genomic variations of 145 genotypes representing seven species (*P. spicata, P. tauri, P. geniculata, P. libanotica, P. strigosa, P. stipifolia, and P. cognata*) from major global lineages, using genotyping-by-sequencing. Our results identify six distinct genetic clusters, with *P. spicata* (a North America species) clearly separated from the other six species underscoring its unique genetic identity. In contrast, the Eurasian species exhibit mixed ancestry, indicating intricate genetic relationships and widespread exchange of genetic material. Furthermore, no single species tree fully captures the relationships among them, implying interactions such as hybridization or gene flows between closely related species. To investigate the evolutional history of Eurasian BBWG species, we reconstructed the species tree topology based on the SNV (single nucleotide variants) matrix, which revealed potential gene flow events. Our findings suggest that the Eurasian BBWG species have undergone reticulate evolution, characterized by substantial gene flow.

## 1 Introduction

Bluebunch wheatgrass (BBWG), (StSt, 2n = 14, or StStStSt, 4n = 28), is a perennial grass belonging to the genus *Pseudoroegneria, Triticeae* tribe. This genus consists of various species including *P. libanotica* (Hack.) D.R.Dewey, *P. strigosa* (Schult.) Á.Löve, *P. stipifolia* (Trautv.) Á.Löve, *P. tauri* (Boiss. & Balansa) Á.Löve, *P. geniculata* (Trin.) Á.Löve, and *P. spicata* (Pursh) Á.Löve among others. This grass has significant ecological and agricultural importance due to its remarkable adaptability to grow under harsh environmental conditions, particularly for the tolerance to drought and soil salinity (Yan and Sun, 2011; Zhai et al., 2024). As climate change continues to exacerbate challenges such as soil salinization, BBWG has emerged as a valuable genetic repository harboring useful diversity for enhancing agricultural resilience in the face of shifting climatic conditions. Efforts to enhance its production and resilience have been undertaken with collaborative initiatives between research institutions and agricultural organizations (Lesica and Atthowe, 2007). Therefore, understanding the scope and distribution of genetic variation among populations will harness genetic diversity for crop improvement and adaptation, and ensure the long-term sustainability of healthy ecosystems.

The St subgenome in BBWG is an ancient and integral component of the *Triticeae* tribe, making significant contributions to biodiversity conservation of the grass family across cool-season regions including Europe, Asia, and North America (Dewey, 1984; Yen and Yang, 2020), covering more than 250 polyploid species (Wang et al., 2020b). The relationship between the St subgenome and other genomes within *Triticeae* has been extensively studied due to its widespread presence. An early study (Wang, 1992) assessing the similarity among basic genomes (E, St, P, N, H, and R) in various diploid hybrids revealed that the E genome showed the strongest relatedness to the St genome. Following the E genome, the P and N genomes demonstrated genetic similarity to the St genome, in descending order. In contrast, the H and R genomes displayed a comparatively distant relationship to the St-E cluster. Moreover, recent studies using molecular approaches further resolved differences between the basic genomes (Cseh et al., 2019; Yu et al., 2008) and identified seven homologous groups of St genome to the A, B, and D subgenomes of common wheat (scientific name at first mentioning), suggesting a conserved cytogenetic collinearity between the St genome and the wheat genome (Wang et al., 2020b).

The divergence of *Pseudoroegneria* species and their genetic relationship have been explored in numerous studies (Yan and Sun, 2011; Gamache and Sun, 2015; Mason-Gamer et al., 2010). These studies suggest that *P. libanotica* underwent the earliest speciation, preserving the most ancient genome among all the *Pseudoroegneria* species. However, most of these studies have focused on single-copy genes, such as the translation elongation factor G (*EF-G*), subunit of RNA polymerase II (*RPB2*), and granule-bound starch synthase I (*GBSSI*) (Yan and Sun, 2011). While phylogenetic analysis based on single copy genes is cost-effective and less prone to the effects of concerted evolution, the limited availability of such genes poses a significant challenge in resolving comprehensive relationships among species. Consequently, results based on single-copy genes often show inconsistencies, leading to controversial relationships (Yan and Sun, 2011; Gamache and Sun, 2015).

Given these inconsistencies in phylogenetic relationships, it is essential to consider additional evolutionary forces, such as gene flow, which can substantially influence allele frequency variation within populations over time (Ellstrand, 2014). To date, gene flow has been documented in many outcrossing grass species, such as ryegrass (*Lolium perenne L.*) (Cunliffe et al., 2004), turtle grass (*Thalassia testudinum*) (Schlueter et al., 1998) and turfgrass (*Poa trivialis L.*) (Brunharo et al., 2024). As an outcrossing lineage, *Pseudoroegneria* species appear to have a higher chance of hybridization or introgression through cross pollination. However, research focusing on the gene flow in *Pseudoroegneria* species remains limited, leaving a gap in our understanding of their evolutionary dynamics.

The genotyping-by-sequencing (GBS) approach has become as a powerful tool for exploring genetic diversity, enabling the detection of extensive single nucleotide variants (SNVs) across the genome. Although GBS has been widely used to study population structure in various species (Niu et al., 2019; Eltaher et al., 2018; Chen et al., 2017), the population structure, genetic diversity, and genetic relatedness of BBWG species have yet to be examined on a whole genome scale. Due to their out-crossing nature, understanding and characterizing the genetic diversity among *Pseudoroegneria* species provides valuable insights for improving breeding efforts to better support ecosystem restoration (Jones et al., 2022). Additionally, GBS enables a comprehensive, whole-genome analysis of population structure, allowing us to uncover patterns of relatedness, genetic differentiation, and gene flow among species. To address existing knowledge gaps, our study leverages the GBS approach to perform a genetic analysis on 145 accessions representing seven *Pseudoroegneria* species collected globally, focusing on the identification of SNVs at the whole genome level. We aim to explore the population structure, genetic diversity and differentiation, as well as to delineate the relationships and gene flow between different *Pseudoroegneria* species.

## 2 Materials and methods

### 2.1 Plant Material

A collection of 145 accessions of *Pseudoroegneria* species were used in this study. All the genotypes were obtained from Genebank at the National Plant Germplasm repository, United States Department of Agriculture (USDA) in Ames, Iowa. The collection consists of 79 accessions of *P. spicata*, 24 accessions of *P. tauri*, 17 accessions of *P. geniculata*, 8 accessions of *P. libanotica*, 7 accessions of *P. strigosa*, 4 accessions of *P. stipifolia*, 1 accession of *P. cognata*, and 5 unclassified accessions (**Supplementary Table S1**). In terms of geographical representation, the collection includes 72 genotypes from the United States, 35 from Iran, 14 from Russia, 8 from Uzbekistan, 7 from Canada, 4 from China, 2 from Ukraine, and 1 each from Mongolia, Kyrgyzstan, and Estonia (**Figure 1**).

**Figure 1.**
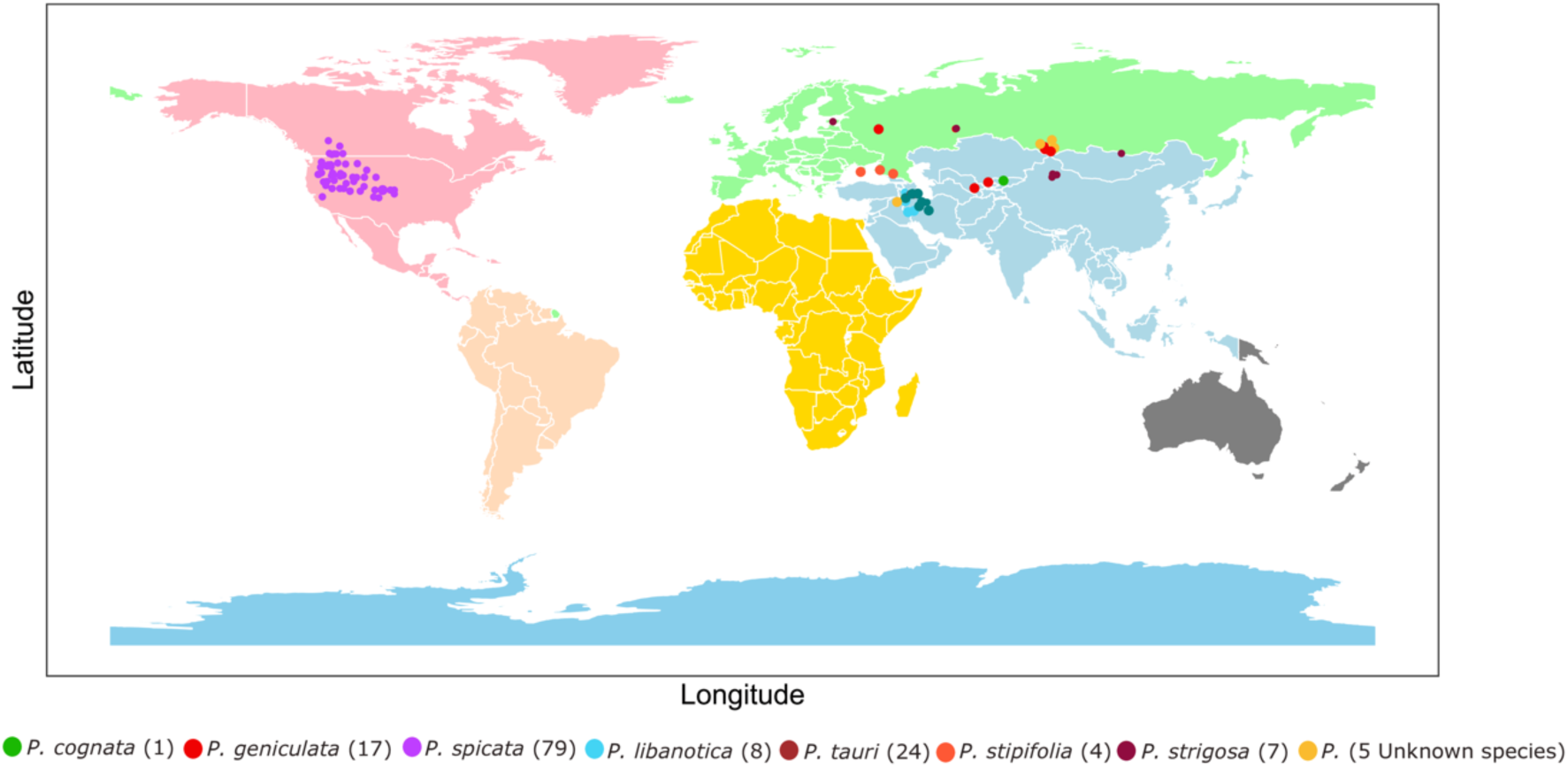
Global map showing the origin collection site of all the *Pseudoroegneria* species sampled in current study. Each dot represents an accession and color coded based on species. The origin sites and species identification are extracted from the USDA Germplasm Resources Information Network.

### 2.2 DNA Extraction and GBS Library Preparation

5-week-old leaf samples were collected for DNA extraction. Leaf tissue was flash frozen with liquid nitrogen and stored at - 80^◦^C prior to DNA extraction. DNA extractions were performed using high-throughput DNA extraction kit (NucleoMag) with a KingFisher 96 instrument (ThermoFisher Scientific). Tissues were disrupted in the lysis buffer and treated with RNaseA to remove RNA contamination. DNA quantification was performed with Quant-iT PicoGreen dsDNA Assay Kit (ThermoFisher Scientific) through fluorescence measured (485nm/535nm, 0.1s) using the Victor XPlate Reader (PerkinElmer). DNA samples were normalized to 20 ng/ul and the processed through an established double digest GBS protocol (Poland et al. 2012).

In brief, 200 ng of DNA from individual samples were digested with the CHG methylation sensitive restriction enzymes *PstI* and *MspI* at 37^◦^C for 2 hrs in a 96 well plate format. After digestion, the DNA were ligated to the sequencing adapters using T4 DNA ligase at 22^◦^C for 2 hrs. The ligation products were deactivated and all 96 samples from each plate were pooled into a single library for 96 samples in a plate. The pooled DNA was used as template for PCR with a short extension time of 30 sec to enrich the library. After PCR, the library was purified using a QIAquick PCR Purification Kit (Qiagen) and quantified by Bioanalyzer (Agilent Technologies) to confirm the fragment size and quality of the library. Sequencing of GBS libraries representing 145 genotypes of *Pseudoroegneria* species were completed on an Illumina NovaSeq 6000 platform (paired-end 150 bp reads).

### 2.3 DNA Sequence Analysis and Variants Calling

An in-house-developed Perl script (https://github.com/USask-BINFO/Bluebunch-wheatgrass-population-structure.git) was used to demultiplex the raw reads. Trimmomatic v.0.38 program was used to trim the adapters, short reads and poor-quality data (Bolger et al., 2014). Leading and trailing bases with quality below 15 and reads shorter than 51 were removed. A total of 1,951 million reads passed filtering and were retained in downstream analysis. Due to the absence of a BBWG reference genome assembly, a tall wheatgrass (*Thinopyrum elongatum*) genome was used as the reference for read mapping due to its close phylogenetic relationship with the species under study (Wang et al., 2020a). This combination enabled more accurate alignment and variant detection making it a suitable reference for comparative genomics and variant analysis. Each sample was aligned to the reference genome using Burrows-Wheeler Aligner (BWA-MEM) v 0.7.18 to generate bam files (Li and Durbin, 2009). Approximately 88% of reads were aligned to the reference genome (Supplementary Table S2). GATK (v4.3.0) variant calling pipeline was used to call SNVs and short INDELs and the creation of VCF files for each sample (Van der Auwera et al., 2013). SNVs variants were extracted from VCF files and filtered with VCFtools (Danecek et al., 2011) with parameters –max-missing 0.4 –maf 0.05 –mac 3 –mQ 20.

### 2.4 Principal component analysis (PCA) and Population Structure Analysis

PCA analysis was performed based on the SNV data using *pca* function of the R package LEA v.2 (Frichot and François, 2015), and the results were plotted with R ggplot2 package (Wickham 2016). All 145 accessions were visualized on the first two principal components. To approximately evaluate the number of populations in *Pseudoroegneria* species, the number of genetic clusters (*k*) within the population was estimated using *cross.entropy* function integrated in non-negative matrix factorization (sNMF) v. 1.2 (Frichot et al., 2014). The optimal *k* value was identified with the lowest entropy value. Population structure and admixture analysis were conducted using LEA and sNMF on the combined dataset of 145 samples representing all seven BBWG species with default settings (**Figure 5**).

### 2.5 Estimation of Heterozygosity (*Ho*), Nucleotide Diversity (*π*), and Tajima’s D (***D***) in Each Species

The observed Heterozygosity (*Ho)* is a measure to describe the proportion of individuals in a population that are heterozygous at a particular genetic locus. It reflects the actual genetic diversity present in the population. *Ho* is calculated based on SNV data as specified in Equation (1), where *A_i_* indicates the individual allele for locus *i*, *f*[*A*_*i*_*A_i_*] is the frequency of homozygote for locus *i, n* is the total number of alleles identified.

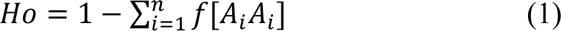

Nucleotide diversity (*π*) is a measure of the genetic variation at the nucleotide level within a population. It quantifies the average number of nucleotide differences per site between two randomly chosen individuals in a population. The calculation of *π* is based on the definition introduced by (Nei and Li, 1979) computing the average number of nucleotide differences per site with adjusted calculations for missing data (Korunes and Samuk, 2021). The expression was shown in Equation (2), where *c_0_ and c_1_* represents the number of two different alleles at a locus, n is the number of individual samples.

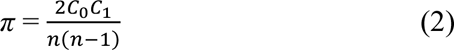

*π* is frequently used as an indicator of the degree of genetic diversity in a population at the sequence level. Tajima’s D is a statistical test used to assess the neutrality of mutations in a population. It compares the number of segregating sites and the average number of nucleotide differences between pairs of sequences in a population to determine if the population is evolving according to neutral evolution or if there are signs of natural selection.

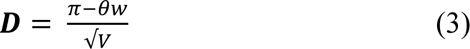

*π* is the pairwise nucleotide diversity. *θw* is the expected number of segregating sites under a neutral model. *V* denote the sampling variance of the difference between the two estimates (*π* and *8θ*). *D* = 0 suggests that population is evolving neutrally. D > 0 reflects that the observed nucleotide diversity is higher than we expect implying that a balancing selection maintains the variants diversity. In contrast, D < 0 suggests a lower diversity than we expected indicating a purifying selection happened in the past.

Nucleotide Diversity and Tajima’s D were calculated for each species using Pixy packages developed by Korunes and Samuk (2021) (https://github.com/ksamuk/pixy) using a 50 kb window across the genome. An in-house-developed python script was used to compute the *Ho* for all the accessions. The script is available in https://github.com/USask-BINFO/Bluebunch-wheatgrass-population-structure.git.

### 2.6 Genetic Differentiation Analysis and Estimation of Pairwise Wright’s Fixation Index (***F*_ST_**)

Wright’s Fixation Index (*F*_ST_) is a measure of population differentiation based on genetic variation. It is calculated as the ratio of the genetic variance between populations or subpopulations over the total genetic variance. When *F*_ST_ =0, it indicates that there is no genetic differentiation between populations. *F*_ST_> 0.25 is considered as large differentiation, suggesting strong genetic separation which may be due to barriers to gene flow, such as geographic isolation or strong selective pressures. We used VCFtools and a 50 kb window across the genome to calculate pairwise *F*_ST_ values. To visualize the *F*_ST_ values as a heatmap (**Figure 6**), we followed a systematic approach. The data was first processed and converted to numeric values, with non-numeric entries replaced by NA. The median value for each species was used to construct the matrix and the data was visualized using the seaborn (Waskom et al., 2021) function in python.

### 2.7 Species Tree Estimation and Gene Flow Analysis

Species-tree inference was performed with the multi-species coalescent model which accounts for the complexities of speciation, incomplete lineage sorting, and gene flow. We use SVDQuartets (Chifman and Kubatko, 2014) implemented in the PAUP software (Swofford, 2003) to construct the species tree. Initially, the filtered SNV dataset was used as input and the VCF format was converted to Nexus format by using vcf2phylip (https://github.com/edgardomortiz/vcf2phylip.git) (Ortiz, 2023). SVDQuartets estimate the topology of the species tree based on the SNV matrix with bootstrapping (200 replications). The gene flow analysis was performed with Dsuite software (https://github.com/millanek/Dsuite.git) (Malinsky et al., 2021). Dtrios command was used to calculate the D statistics and the *f*4-ratio for all combinations of trios of species. The *f*-branch metric was calculated with subcommand Fbranch indicating the gene flow between specific branches. *f*-branch values were plotted using a python script (dtools.py) in the same package of Dsuite (**Figure 7**).

## 3 Results

### 3.1 Genome-wide Discovery of Genetic Variants and Genetic Diversity in *Pseudoroegneria* Species

Due to the close genetic relationship between *Thinopyrum elongatum* and *Pseudoroegneria*, the chromosome-level assembled genome of *Thinopyrum elongatum* was used as a reference for mapping GBS reads. A total of 1.95 billion clean reads were obtained from 145 lines after filtration, with an average mapping rate of 88% to the reference (**Supplementary Table S1**). A total of 31,489,307 genetic variants were identified, including 3,186,094 INDELs and 28,277,996 SNVs. Among all the SNVs, 99.41% are located on the seven chromosomes, with the remainder on unannotated scaffolds (**Supplementary Table S2**). The initial VCF file was subjected to stringent filtering criteria (see Methods), which reduced the number of SNVs to 59,511. These filtering thresholds ensure that the dataset retain high-confidence, moderately frequent variants with adequate coverage, thereby enhancing the reliability and robustness of downstream analyses.

The ratio of transition and transversion type of SNVs (Ts/Tv) is 1.65, with 33.1% of A/G and 33.0% of C/T transitions, and a total of 40.1% of all four types of transversion (A/C, A/T, C/G, and G/T) (**Supplementary Table S3**). The average density of filtered SNVs across all chromosomes was 13 SNVs/Mb, with chromosome 4 exhibiting the lowest density (9.5 SNVs/Mb) (**Figure 2; Supplementary Table S2**). The highest density was observed on chromosome 5, followed by chromosome 1 after filtration (**Supplementary Table S2**) suggesting a relatively high abundance of diversity on these two chromosomes.

**Figure 2.**
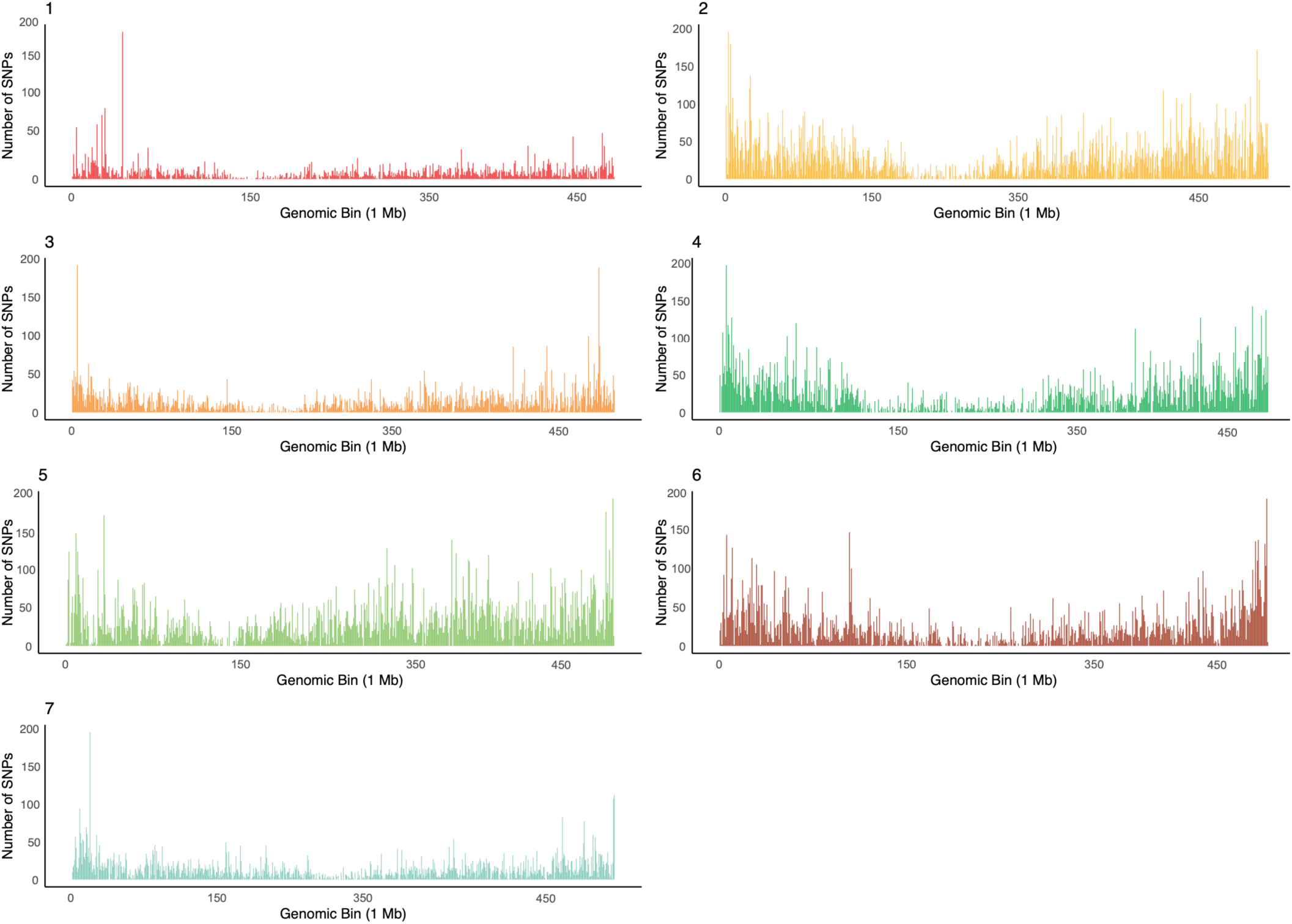
SNV density across chromosomes of *Pseudoroegneria* species. Each plot represents the SNV density distribution (post-filtration) for one chromosome, with chromosome numbers labeled at the top left of each panel. The x-axis shows genomic bins of 1 Mb, and the y-axis indicates the number of SNVs within each bin. The variation in SNV density across the genome reflects the genomic diversity in *Pseudoroegneria* species.

### 3.2 Observed Heterozygosity (*Ho*), Nucleotide Diversity (π) and Tajima’s D for Individual Species

To investigate the evolutionary dynamics of various *Pseudoroegneria* species, we focused on three key metrics including the observed *Ho,* π, and Tajima’s D within each species. The biological significance of these metrics is detailed in the Methods section. Analysis of these metrics would offer a comprehensive view of the evolutionary process shaping these species, such as selection pressure, historical demographic events.

The results revealed significant variation among populations of different species (**Figure 3**). The *P. spicata* population has substantially lower π and *Ho* compared to other populations, coupled with a negative Tajima’s D value. This suggests a limited pool of genetic variation, possibly due to historical bottlenecks or founder effects, where the population was established with a small number of individuals and has experienced a recent population expansion. In contrast, the *P. stipifolia* population displayed the highest genetic diversity, *Ho*, and Tajima’s D values implying a high degree of variation within the population and likely resulting from stable population dynamics with ongoing balanced selection that maintains a wide array of genetic variation. Other than these two extreme cases, the other four species displayed increased π and Tajima’s D value in the order of *P. tauri*, *P. libanotica*, *P. strigosa*, *P. geniculata*. However, the observed *Ho* displays a slightly different pattern among these four species, with *P. geniculata* and *P. strigosa* exhibiting the next highest *Ho* values. Additionally, the *Ho* values of *P. stipifolia* and *P. strigosa* span a wider range within their populations compared to the other species suggesting more variations across the individual accessions.

**Figure 3.**
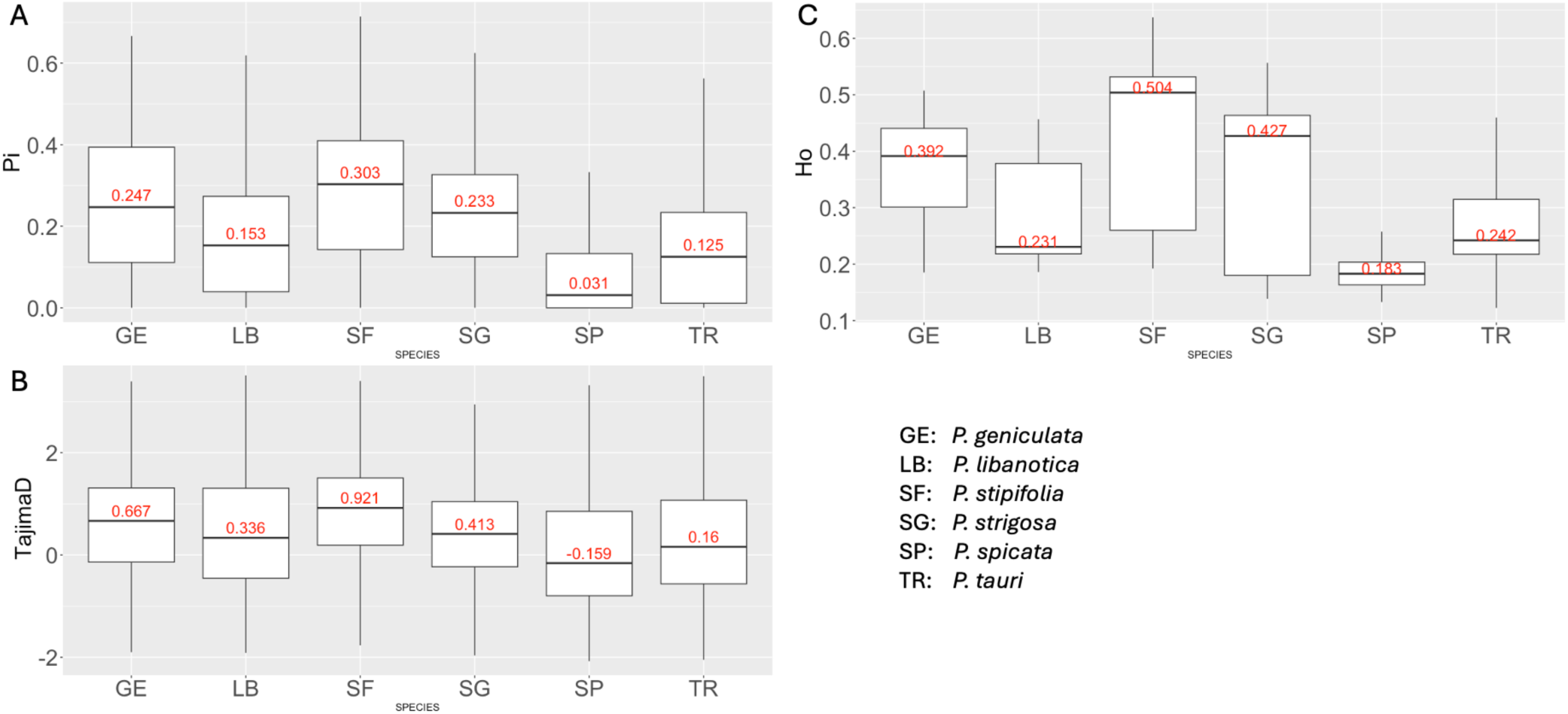
(A and B), Nucleotide diversity per base pair (π) and Tajima’s D estimation within *Pseudoroegneria* species across the genome in 50 kbp windows. (C) The observed heterozygosity (*Ho*) distribution within *Pseudoroegneria* species. Boxplots depict the median (center bar with values in red) with the outliers removed for clarity. (Species abbreviations, GE: *P. geniculata*, LB: *P. libanotica*, SF: *P. stipifolia*, SG: *P. strigosa*, SP: *P. spicata*, TR: *P. tauri*).

### 3.3 Population Structure and Genetic Relationship among *Pseudoroegneria* Species

Population structure analysis of the 145 BBWG accessions was carried out to better understand the effectiveness of the detected SNVs in distinguishing closely related species and the extent of allele sharing among subspecies. Three approaches were employed based on 59,511 SNVs (post-stringent filtration) including PCA analysis, admixture inference, and phylogenetic tree reconstruction. The PCA results revealed that all individuals were clustered into six distinct groups (**Figure 4A**). The first two principal components accounted for 14.35% and 6.88% of total genetic variation, respectively, clearly separating the *P. spicata* (in purple). Interestingly, *P. tauri* accessions (in teal) were clustered into two sub-populations, with each group mixed with individuals from other species. The remaining clusters contained a few individuals from each species, suggesting closer genetic relationships or potential hybridization events between these species. The clustering pattern observed in the PCA highlights the genetic distinctiveness of *P. spicata*. In addition, the UPGMA phylogenetic tree based on the same set of SNVs supports the clustering observed in the PCA, with all the *P. spicata* accessions clustering together forming a distinct clade. While the other five species exhibit interbranching except *P. geniculata* showing most of the individuals in the same branch (**Figure 4B**).

**Figure 4.**
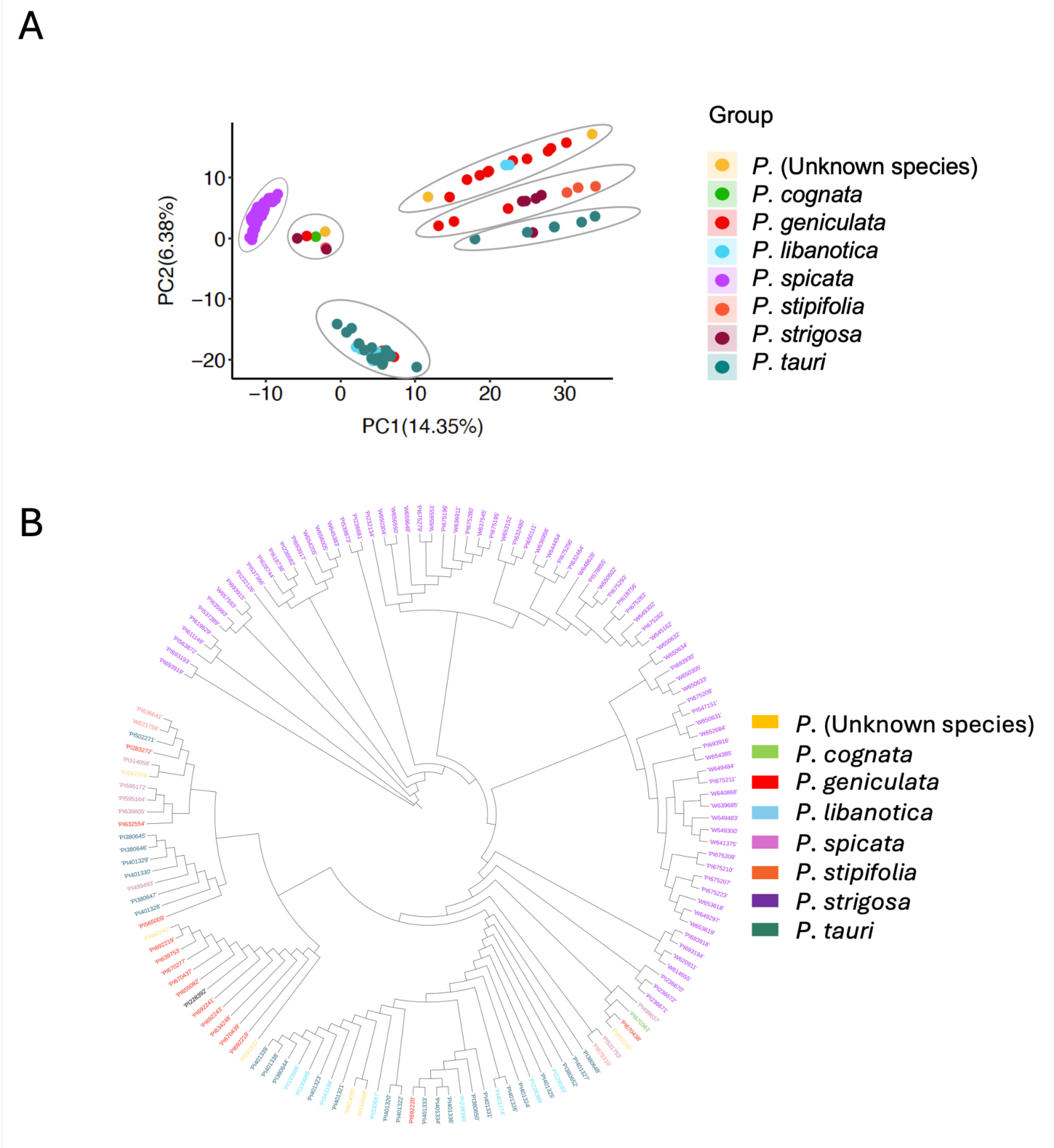
Principal coordinate analysis (A) and Phylogenetic dendrogram (B) of 145 accessions from seven *Pseudoroegneria* species. Individuals are color coded based on the taxonomic identification. The species identification was extracted from the USDA Germplasm Resources Information Network.

Furthermore, we assessed the population structure using the sNMF package (Frichot et al., 2014). To define the optimal number of genetic clusters (*k*) for admixture inference, we explored values from one to ten, with 100 replicates for each *k*. The lowest cross-entropy value (*k*=4) was identified as the best *k* (**Figure 5A**) (Frichot et al., 2014). Therefore, we utilized a range of *k* values (2-4) in subsequent analyses, as shown in **Figure 5B**. The sNMF model-based structure analysis with *k* = 2 clustered the *Pseudoroegneria* accessions into two main ancestral populations, with *P. spicata* (purple bar covered) being clearly distinct and 100% from one of the ancestors, while the other species displayed mixed ancestry predominantly in yellow. This result suggested that the Eurasian accessions can be divided into two ancestral populations while *P. spicata* is likely derived from one of the ancestors in Eurasia.

**Figure 5.**
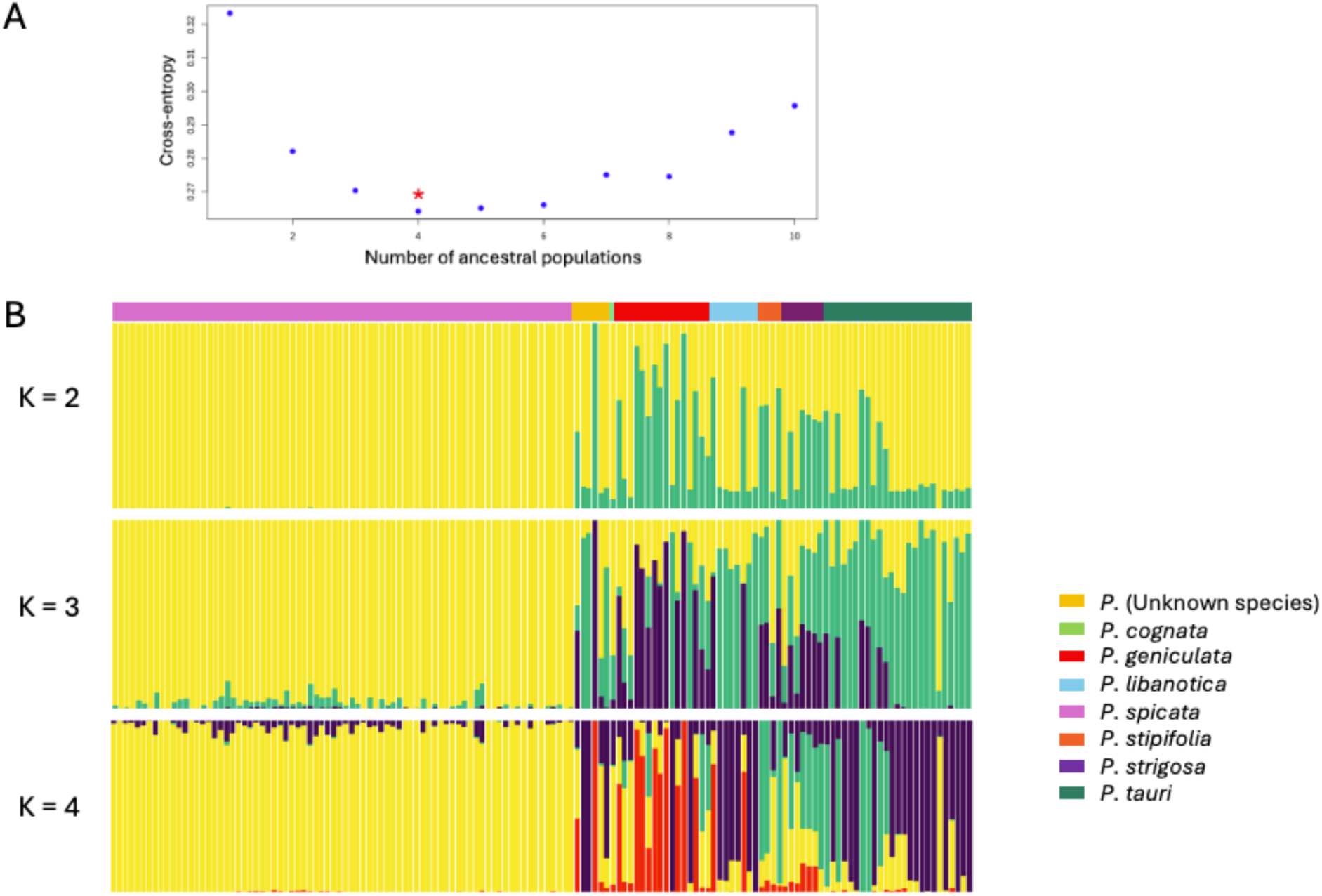
(A) *k* value estimation using cross entropy. (B) Population structure for *Pseudoroegneria* species. Each bar represents an accession. The chart was plotted based on different *k* values (*k* = 2 to 4). The bar on the top showing different colors represents the species as annotated.

As the value of *k* increased to three, the Eurasian accessions continued to separate into three subgroups. At *k* = 4, a few accessions clustered into four mixed ancestry groups, while most still maintained two or three mixed ancestry components. In contrast, *P. spicata* exhibited a very low percentage of mixed ancestry even as the *k* value increased. Furthermore, the Q-value for a cluster represents the fraction of an individual’s genome inherited from that cluster, reflecting its association with the ancestral group. For Eurasian species, a high Q-value (e.g., >80%) indicates a predominantly association with a single ancestral cluster. According to the Q-value results (**Supplementary Table S4**), 5 out of 17 accessions in *P. geniculate*, 5 out of 8 in *P. libanotica*, 3 out of 7 in *P. stipifolia* and 18 out of 24 in *P. tauri* show strong associations with different ancestry, rather than admixture. These accessions, with their high Q-values to specific ancestral group can serve as valuable pure lines for future research and breeding programs.

### 3.4 Population Differentiation

Pairwise genetic differentiation (*F*_ST_) between six major species was computed genome-wide to examine the divergence of each subspecies. The differentiation between the Eurasian BBWG subspecies shows relatively low *F*_ST_ values (0.01 - 0.066) (**Figure 6**), indicating a close genetic relationship and suggesting that these populations have maintained significant gene flow or have diverged relatively recently from a common ancestor. This close genetic affinity among Eurasian subspecies is not surprising due to their geographical proximity and the potential interbreeding.

**Figure 6.**
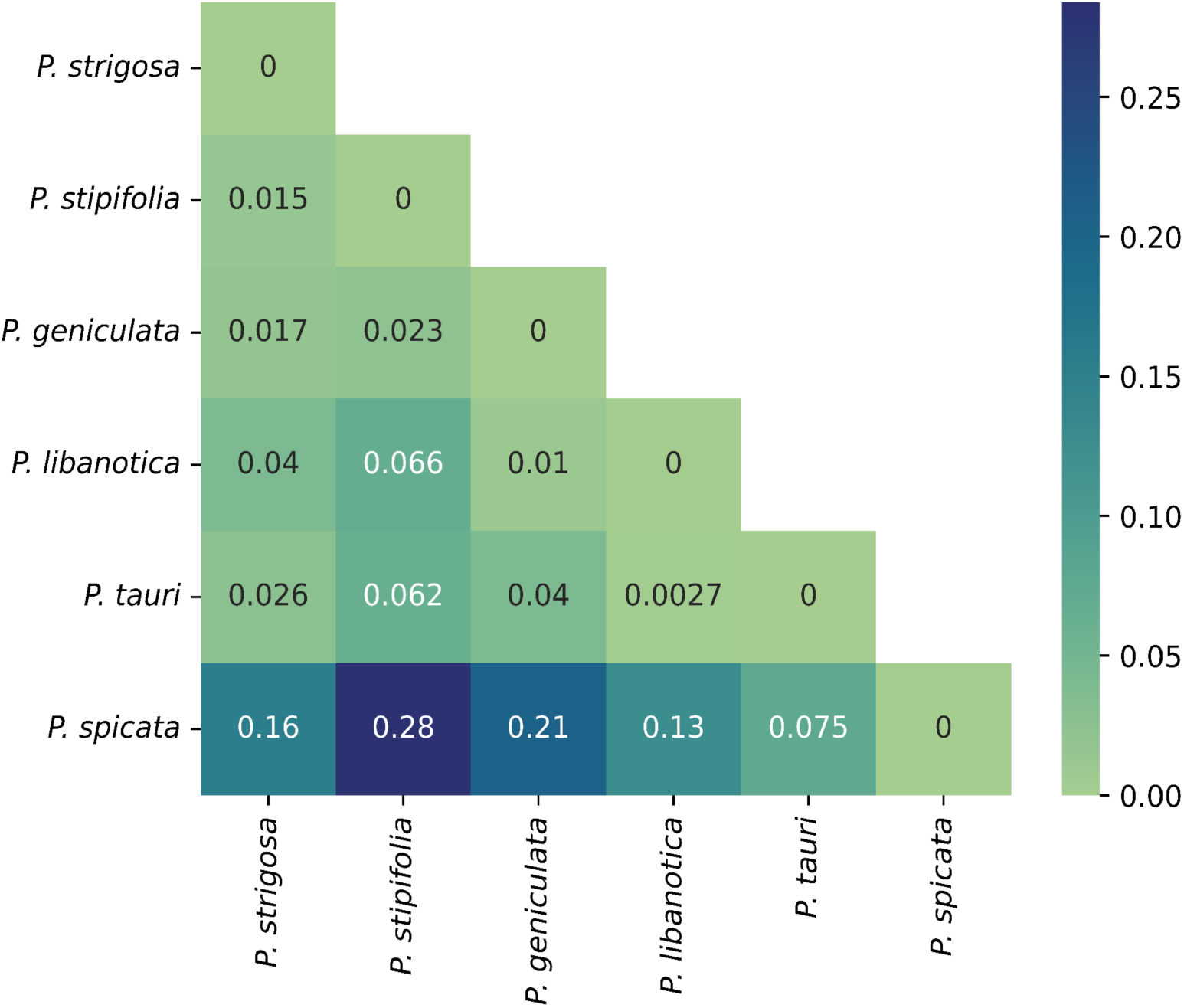
**Heatmap of median *F*_ST_ value for pairwise comparison of *Pseudoroegneria* species (window size = 50 kbp).**

In contrast, *P. spicata*, which is native to North America, exhibits significantly higher *F*_ST_ values when compared to most of the Eurasian species (0.13 - 0.28) indicating substantial genetic divergence between them. However, the *F_ST_* value (0.075) between *P. tauri* and *P. spicata* was significantly lower than the rest of Eurasian species (**Figure 6**), suggesting a somewhat closer genetic relationship between them, possibly due to historical gene flow or shared ancestral polymorphisms that have been maintained in both populations. Interestingly, examining the *F*_ST_ values across individual chromosomes reveals that most chromosomes exhibit a consistent pattern of genetic differentiation. However, chromosome 4 exhibits a higher *F*_ST_ between *P. spicata* and *P. geniculata* (**Supplementary Figure S1**).

### 3.5 Species-tree Inference and Gene-flow Estimation

Since the population structure analysis and individual phylogenetic tree do not adequately represent a comprehensive and precise representation of the relationships between Eurasian species, delineating species boundaries within these populations remains a significant challenge. We therefore inferred a species tree using 2-4 accessions from each species to obtain an initial picture for exploring the evidence of hybridization or gene flow events (Accession list in **Supplementary data Table S5**). A quartets-based method (SVDQuartets) (Chifman and Kubatko, 2014) was employed to infer the species tree with rice as an outgroup species (**Figure 7A**). This tree provides strong confidence (>98%) in the close relationship between *P. spicata* and a clade formed by *P. libanotica* and *P. tauri*, indicating a well-resolved phylogeny for these three species. In contrast, the relationship among *P. strigosa*, *P. geniculata*, and *P. stipifolia* are supported with relatively moderate confidence.

**Figure 7.**
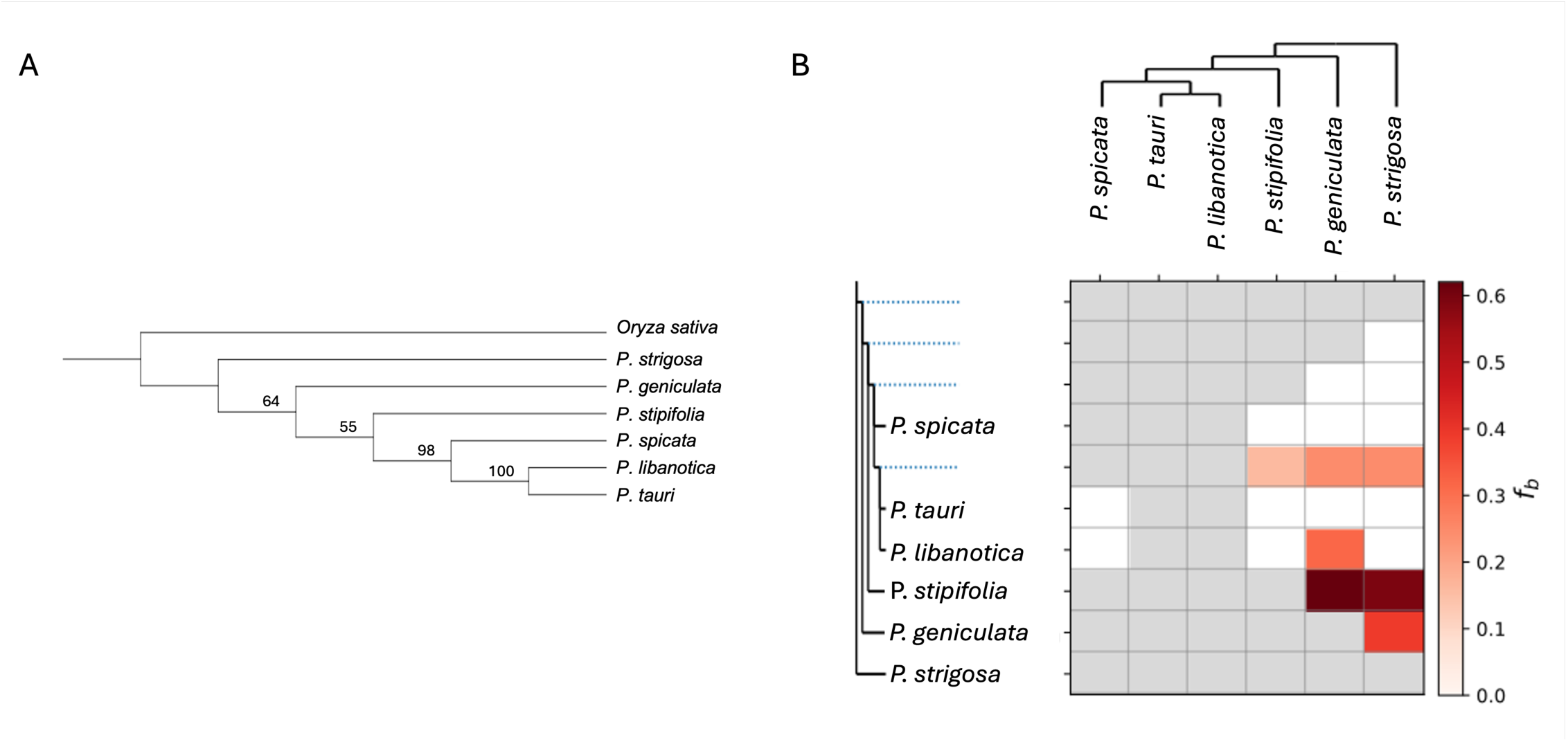
(A) Species tree inference based on quartet relationships (SVDQuartets). The number of replications for bootstrap is 200. (B) The f-branch statistics identify excess sharing of derived alleles between species (labelled on the x-axis) and branches (labelled on the y-axis). The tree displayed on the x-axis is the reference tree inferred using SVDQuartets, the expanded tree along the y-axis displays all the possible branches, the branches containing more than two leaves were not displayed in the representation.

Next, we used this inferred species tree as a reference tree and calculated *f*-branch statistics to assign evidence of gene flow to specific branches based on correlated *f*_4_-ratio tests (Malinsky et al., 2021) using the *f*-branch metric introduced by Malinsky et al., 2018. Given a specific tree either true or hypothesized, the *f*-branch statistics reflect excess sharing of alleles between the populations or species to other branches. As *f* statistic values for different combinations of ((A,B),C) groups are not independent due to the branch sharing, this method aimed to obtain branch-specific estimates of excess allele sharing that are expected to be less correlated (Malinsky et al., 2018). The top two *f*-branch values (62.04% and 58.25%) are observed in the pairs of *P. stipifolia*/*P. geniculata* and *P. stipifolia*/*P. strigosa*, respectively, indicating substantial sharing of excess alleles between each pair and possibly hybridization between each pair (**Figure 7B**). Signals for the gene flow between *P. libanotica* to *P. geniculata* and between *P. geniculata* to *P. strigosa* are less pronounced (30% - 40%). In addition, the *f* branch signals between each pair are only observed in one direction suggesting a single-directionality introgression (Malinsky et al., 2018).

## 4 Discussion

### 4.1 Genetic Relationships Between St and E Genomes

Over the past few decades, significant progress has been made in resolving phylogenetic relationships within the *Triticeae* tribe using cytogenetic and molecular evidence. However, some aspects of these phylogenies remain incomplete and unclear, particularly concerning the relationships between basic genomes (Hsiao et al., 1995; Wei and Wang, 1995; Mason-Gamer et al., 2002; Yu et al., 2008). Several cytogenetic and molecular studies have suggested a close relationship between the St and E genomes (Wang, 1989; Bieniek et al., 2015), yet they have not fully elucidated the extend of this relatedness.

Recent advancements have provided assembled genomes for both the St and E genomes, revealing substantial differences in genome size. The St genome from *P. libanotica* has an assembled size of 2.99 Gb, with an estimated total size of 3.1 Gb (Zhai et al., 2024). In contrast, the E genome from *T. elongatum* comprises 4.58 Gb, representing for 95.8% of its estimated genome size of 4.78 Gb (Wang et al., 2020a). Despite both genomes being published, at time of submission, the reference of St genome was not publicly available, presenting challenges in accurately characterizing its genetic relatedness to other subgenomes. Given the availability and completeness of E genome, our study addresses this gap by mapping the GBS reads from all *Pseudoroegneria* accessions to the E genome. The mapping rate ranged from 85% to 91%, with an average of 88%, indicating a notable level of similarity between the St and E genomes.

### 4.2 Origin of BBWG

Due to the nature of outcrossing and polyploidy, *Pseudoroegneria* species rely on cross-pollination for reproduction, exhibiting complex evolutionary histories shaped by hybridization, introgression, and extensive gene flows. This complexity requires extensive effort to disentangle their genomic structure and evolutionary dynamics. Previous studies have suggested that *P. libanotica* represents an early-diverging lineage within the genus (Zhai et al., 2024). In this study, we expanded upon these findings by analyzing a broader range of populations, encompassing seven major lineages. Our results indicate that *P. tauri* and *P. libanotica* occupy similar position in the phylogenetic tree and share comparable ancestral genetic components. This strong genetic affinity suggests that both species may have diverged early in the evolutionary history of *Pseudoroegneria.* The close relationship between *P. tauri* and *P. libanotica* raises the possibility that they share a common progenitor, which could either be missing from our collection or possibly extinct.

### 4.3 Genetic Variation and Evolutionary Insights in *Pseudoroegneria* Species

Genetic variation is a cornerstone of evolution, providing the diversity necessary for populations to adapt to natural selection and genetic drift. This study presents a comparative analysis of genetic diversity among *Pseudoroegneria* species using key metrics such as nucleotide diversity (π), Tajima’s D, and observed heterozygoisty (*Ho)*. Notably, *P. stipifolia* emerges as the most genetically diverse species, while *P. spicata* exhibited the lowest genetic diversity, despite having the largest population sample in this study.

A previous study (Larson et al., 2000) analyzed the diversity of different cultivars using amplified fragment length polymorphism (AFLP) loci and compared the genomic variation to wild populations, including a collection of 25 wild *P. spicata* lines. Their findings suggested that genetic diversity has not been significantly reduced by genetic drift since the divergence of the cultivars. Our results, based on Tajima’s D, indicate that *P. spicata* likely originated from a small number of individuals and has undergone recent population expansion in western North America. This aligns with the observed minimal differences in genetic diversity between cultivars and their wild counterparts. Together, these findings suggest that genetic drift has had limited impact on *P. spicata*’s diversity over time, and the species has maintained remarkable genetic stability, underscoring the resilience of its genetic makeup.

However, despite this genetic stability, there is a pressing need to enhance genetic diversity within *P. spicata*. Incorporating genetic material from other species into breeding programs could bolster resilience and ensure long-term sustainability.

In terms of genetic diversity across all the *Pseudoroegneria* species, the variation in SNV density across the chromosomes points to underlying biological significance. The elevated SNV density observed on chromosomes 5 and 1 indicates regions of increased genetic diversity, potentially identifying evolutionary hotspots within the *Thinopyrum elongatum* genome. These regions may be influenced by natural selection or exhibit higher rates of recombination, contributing to the accumulation of genetic variation. In contrast, the reduced SNV density on chromosome 4 suggests a more conserved genomic region, likely due to the presence of essential genes where variation is less tolerated.

More interestingly, most of the chromosomes in *P. spicata* show a consistent level of differentiation compared to Eurasian species. However, chromosome 4 stands out, showing a unique pattern of differentiation between *P. spicata* and *P. geniculate* (**Supplementary Figure S1**). This anomaly suggests that chromosome 4 may harbor specific regions or loci under distinct selective pressures, potentially due to local adaptation, genetic drift, or historical events unique to this chromosome. The distinct genetic landscape of chromosome 4 highlights its potential role in driving adaptive differences or reflecting unique evolutionary histories not observed in other chromosomes.

This study underscores the importance of genetic diversity in evolutionary processes and provides new insights into the genetic dynamics of *Pseudoroegneria* species, particularly the unique evolutionary trajectory of *P. spicata* and the exceptional characteristics of chromosome 4.

### 4.4 Gene Flow, Hybridization and Evolutinoary Dynamics in Eurasian BBWG Species

Gene flow, the transfer of genetic material between populations, plays a vital role in the genetic diversity and adaptability of species (Ellstrand 2014). Based on the results of population structure, PCA, and phylogenetic analyses (**Figures 5 and 6**), our findings suggest that the presence of shared genetic markers among Eurasian species, particularly for *P. geniculata*, *P. stipifolia*, and *P. strigose.* This genetic overlap complicates the delineation of clear species boundaries, indicating that these species have not fully diverged genetically. Instead, they still maintain a continuum of genetic exchange, presenting a considerable challenge for phylogenetic analyses.

A hybridization event was previously proposed in a study examining the genome constitutions of *P. geniculata* (PI634248) and its subspecies *P. geniculata* spp. *scythica* (PI502271) and *P. geniculata* ssp. *pruinifera* (PI547374) (Yu et al., 2010). This study revealed that *P. geniculata* contains the St genome, which is similar to *P. libanotica*, while its two subspecies share the St genome with *P. stipifolia.* Additionally, some species exhibit partial similarity with the E genome, suggesting hybridization between the St and E genomes. Notably, two accessions from this study were also included in our *P. geniculate* collection. The concordance between our findings and the previous study underscores the significant role of hybridization in shaping the genetic landscape of these species.

The relationships among diploid species within the Triticeae tribe are often complicated by intergeneric and interspecific hybridizations, introgression events, and the incomplete lineage sorting of ancestral polymorphisms (Mason-Gamer, 2005; Kellogg et al., 1996; Escobar et al., 2011). Despite the challenges posed by this reticulate pattern of evolution, our study identifies a high-confidence species tree for the relationships among *P. spicata*, *P. tauri*, and *P. libanotica*. This close relationship is further supported by *F*_ST_ values and results from phylogenetic analyses of the *RPB2* gene sequences, which group these three species into a distinct clade (Yan and Sun, 2011). Interestingly, the results from the same study separated *P. spicata* into several groups, a pattern not reflected in our population structure analysis. This discrepancy may rise from two factors: (1) our seed collection of *P. spicata* might be an underrepresentation of its genetic diversity due to a rapid population expansion, and (2) the evolutionary markers in *P. spicata* may be more conserved within the species rather than between species. This internal homogeneity could mask broader phylogenetic signals, leading to differences between our findings and previous analyses.

In conclusion, this study provides a comprehensive overview of the genetic diversity and evolutionary dynamics within *Pseudoroegneria* species, revealing critical insights into their genetic structure and species relationships. We found high genetic homology between the St and E genomes, indicating a deep evolutionary connection despite their size differences. *Pseudoroegneria spicata* shows reduced genetic diversity, likely due to a recent population expansion, while *P. stipifolia* maintains a high genetic diversity and stability. The substantial gene flow among Eurasian BBWG species, as evidenced by results of population structure analysis and gene flow estimation, complicates species differentiation. However, our analysis confirms strong phylogenetic relationships among key species. Overall, these findings enhance our understanding of *Pseudoroegneria* genetic diversity and evolution, offering valuable information for future breeding and conservation strategies.

## Author contribution statement

YJ: Conceptualization, Data curation, Formal analysis, Investigation, Writing-original draft, NK: Data curation and Analysis, Draft editing, RC: Methodology, SP: Resources, Methodology, ZW: Data curation, PH: Resources, BB: Resources, AS: Conceptualization, Supervision, Funding acquisition, LJ: Conceptualization, Supervision, Funding acquisition.

## Acknowledgements

We thank the ADF under the Ministry of Agriculture of Saskatchewan, the Saskatchewan Cattleman’s Association and NSERC for their funding support. We also extend our gratitude to the Germplasm Resources Information Network (GRIN) for providing the seeds material essential for our study.

## Appendix A. Supplementary data

Supplementary data to this article can be found online.

## Data availability

Raw GBS sequencing read data are available on the Sequence Read Archive (SRA) under BioProject PRJNA1228636 and project accession PRJEB86311. Sample information is documented in Supplementary Table S1. In-house-develop script can be found on Github (https://github.com/USask-BINFO/Bluebunch-wheatgrass-population-structure.git).

## Supplementary tables

**Table S1** Origin, ploidy and read mapping rate for each accession.

**Table S2** SNV frequency for each chromosome.

**Table S3** The ratio of transitions and transversions.

**Table S4** Q-value for each accession from population structure analysis (k = 4).

**Table S5** Accessions selected for inference of the species tree.

## Supplementary Figure

**Supplementary Figure S1.**
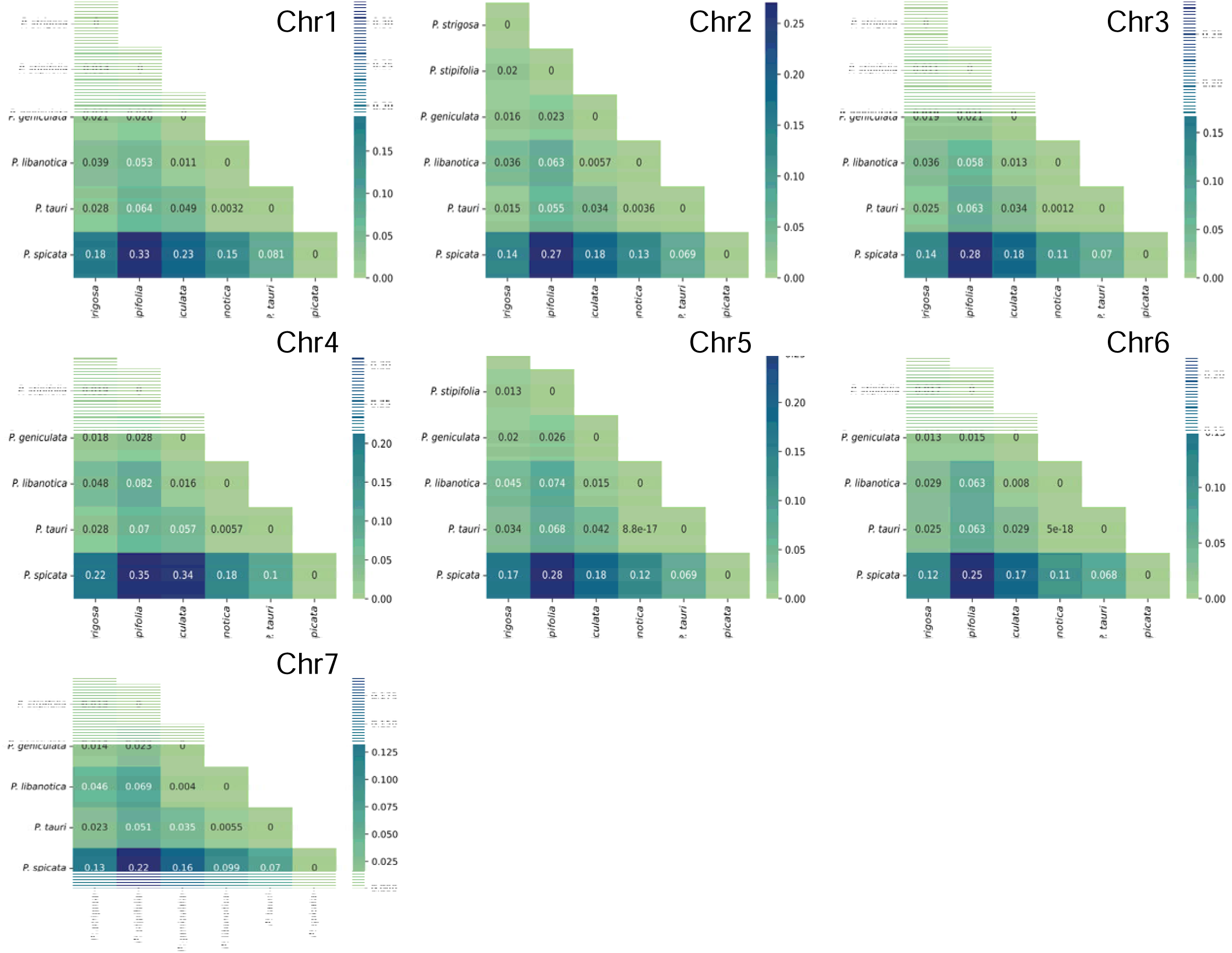
Heatmap of median *F*st value for pairwise comparison of *Pseudoroegneria* species (window size = 50 kbp) for eac chromosome.

